# *In Silico* Identification of Three Types of Integrative and Conjugative Elements (ICEs) in *Elizabethkingia anophelis* Strains Isolated from Around the World

**DOI:** 10.1101/402107

**Authors:** Jiannong Xu, Dong Pei, Ainsley Nicholson, Yuhao Lan, Qing Xia

## Abstract

*Elizabethkingia anophelis* is an emerging global multidrug-resistant opportunistic pathogen. We assessed the diversity among 13 complete genomes and 23 draft genomes of *E. anophelis* derived from various environmental settings and human infections from different geographic regions around the world over past decades from 1950s. Thirty-one of these 36 strains harbor integrative and conjugative elements (ICEs). A total of 52 ICEs were identified, and categorized into three ICE types based on the architecture of signature genes in the conjugation module. The type II and III ICEs were found to integrate into regions adjacent to tRNA genes, while type I ICEs used a variety of integration sites, inserting into intergenic regions or even directly into a gene, sometimes disrupting gene function. Integrases such as tyrosine recombinases, serine recombinases and DDE transposases were found in most ICEs. The ICEs carry various cargo genes including transcription regulators and those involved in antibiotic resistance. The CRISPR-Cas system was found in nine strains, including four strains in which CRISPR-Cas machinery and ICEs co-exist. ICE distribution in the strains showed no geographic or temporal patterns. The ICEs in *E. anophelis* differ in gene structure and sequence from CTnDOT, a well-studied ICE prevalent in *Bacteroides* spp. This is the first set of ICEs identified in the family Flavobacteriaceae. As a prevalent type of mobile genetic elements in various strains of *E. anophelis* around the world, the categorization of ICEs will facilitate further investigations such as virulence, genome epidemiology and adaptation genomics of *E. anophelis.*

**Importance:** *Elizabethkingia anophelis* is an opportunistic human pathogen, and the genetic diversity between strains from around the world becomes apparent as more genomes are sequenced. The Integrative Conjugative Element (ICE), found in many bacterial species, contains genes for transfer via conjugation and integration into the chromosome, along with various cargo genes. ICEs are identified in 31 of 36 strains and categorized into three types based on architecture of modular genes, integrases, and integration sites. ICE distribution in different strains displays no spatial and temporal patterns. Several ICE-containing strains also possessed CRISPR-Cas units, considered to be the bacterial adaptive immune system providing protection against phage and predatory mobile genetic elements. This co-existence suggests that ICEs are beneficial or at least not harmful to the bacterial cells they inhabit. ICEs as a component of the mobile genetic repertoire enable recipients to resist antibiotics, survive disinfecting agents, and adapt to various ecological niches.

## Introduction

The genus *Elizabethkingia* belongs to family *Flavobacteriaceae*, and was recognized as distinct from the genus *Chryseobacterium* in 2005 (1). The two species initially recognized were *E. meningoseptica*, named based on its initial isolation as the causative agent for neonatal meningitis (2), and *E. miricola*, an isolate obtained from condensation water of Space Station Mir (3). A third species, *E. anophelis,* was proposed in 2011(4) based on the description of the type strain R26^T^ that was originally isolated from the midgut of *Anopheles gambiae* mosquitoes maintained in Stockholm University in Sweden (5). In 2015, a new species, *E. endophytica* was proposed (6) but whole genome sequence (WGS) based genome comparison revealed it to be a homotypic synonym of *E. anophelis* (7, 8). Additional species have since been added to the genus (8).

In 2011, the first human infection attributed to *E. anophelis* was documented, a neonatal meningitis that occurred in Central Africa Republic (9). Later, an *E. anophelis* outbreak in an intensive-care unit in Singapore was reported (10), followed by the worrying account of *E. anophelis* transmission from a mother to her infant (11). In 2016, a large *E. anophelis* outbreak occurring in Wisconsin, USA, was unusual in that a substantial proportion of patients were not already hospitalized, and were instead admitted directly from their homes. No indication of human-to-human transmission was found (12-14). Human cases of *E. anophelis* have been reported with increasing frequency around the world (13, 15-17), aided in part by improved identification methods, and several strains previously described as *E. meningoseptica* were determined to actually belong to the *E. anophelis* species (18). In addition to human-derived strains, multiple strains of the species have been isolated from its namesake, the mosquito, and their genomes have been sequenced. Both the type strain R26^T^ and strain Ag1 were derived from *An. gambiae* mosquitoes (19, 20). Strain EaAs1 was isolated from *An. stephensi* mosquitoes (21), and strains AR4-6 and AR6-8 were obtained from wild caught specimens of the mosquito *An. sinensis* in China (20). There has been no epidemiological link between human cases and mosquitoes, and the circumstances of certain outbreaks preclude any possibility of mosquitos being a vector for *E. anophelis.*

Genome comparison of pathogenic bacteria has greatly increased our understanding of the evolution, pathogenesis and epidemiology in many pathogen outbreak investigations (22-26). Horizontal transfer of mobile genetic elements between bacterial strains occurs continually, and is a major source of genetic variation in bacteria (27). One group of modular mobile genetic elements, known as integrative and conjugative elements (ICEs), are capable of transferring between bacteria horizontally via conjugation (28-30). ICEs integrate into a host chromosome and replicate along with the genome (28, 31). The cargo genes that are brought in by the ICEs may endow the recipient bacteria with new phenotypes (32). For example, in *Pseudomonas syringae* pv. *actinidiae*, resistance to copper, arsenic and cadmium was attributed to the resistant genes that were carried in ICEs (33). In *Helicobacter pylori*, ICE *tfs4* has been identified in certain strains mediating gastric disease. The presence of *tfs4* ICE has been associated with virulence, although mechanisms behind the virulence phenotype remain unknown (34).

Genome analysis of various *E. anophelis* strains has revealed substantial genetic diversity (11, 16, 35-39). The ICE named ICE*Ea*1 has been identified in the strains associated with the outbreak in Wisconsin, and unrelated strains from the outbreak in Singapore (40). In this study, we searched for ICEs in 13 complete and 23 draft genomes of *E. anophelis* strains around the world by locating clusters of transfer (*tra*) genes. Based on the architecture of conjugation modules and associated signature genes, three types of ICEs were recognized. As *E. anophelis* is recognized as an environmental bacterium, the high prevalence and diversity of ICEs as well as the mosaic integration pattern in different strains suggest that ICEs play a significant role in shaping its genome to adapt to different environmental niches. In recent years, whole-genome sequencing has been utilized in epidemiological studies to type outbreak strains, acquire genomic determinants of virulence and antibiotic resistance in outbreaks, and estimate pan-genome diversity (41-44). The identification and classification of ICEs is expected to facilitate the genomic epidemiological studies of the pathogenic strains of *E. anophelis* in the future.

## Results

### Origin and geographic distribution of the strains

In this study, 13 complete genomes and 23 draft genomes of *E. anophelis* were compared. Mosquito-derived strains included R26^T^ and Ag1 from *An. gambiae* (19, 20), EaAs1 from *An. stephensi* (21) and AR4-6 and AR6-8 from wild caught *An. sinensis* (20). Strains that were associated with human infections and recent outbreaks were from the Central Africa Republic (9, 39), Singapore (35), Hong Kong (11), China (45), Wisconsin, USA (13, 14), and Taiwan (16). Geographic distribution of included isolates was spread across Asia, Europe, Africa and North America (Figure 1). The available metadata for the strains is noted in Table 1, along with the WGS accession numbers.

**Table 1.** *Elizabethkingia anophelis* strains in the study

| WGS Accession No. | Level | Lineage | Strain | Source | Region | Collection Time | ICE type (n)<sup>a</sup> | CRISPR (n)<sup>b</sup> |
| --- | --- | --- | --- | --- | --- | --- | --- | --- |
| ERS1197909<sup>c</sup> | Draft | II | AmMS250/CIP104057 | Human patient | US | 1994 | II (1) | No |
| CP023010.1 | Complete | II | FDAARGOS-198 | Human patient | Sweden | Unknown | I (2) | No |
| MAHA01 | Draft | II | CSID 3000516074 | Human patient | Illinois, US | 2016 | I (1) | No |
| CP016373.1 | Complete | II | 3375 | Human patient | South Carolina, US | 1957 | I (1) | No |
| MAHS01 | Draft | II | E6809 | Human patient | California, US | 1979 | II (1) | No |
| CP014340.1 | Complete | II | F3543 | Human patient | Florida, US | 1982 | II (1) | No |
| CP016370.1 | Complete | II | 0422 | Human patient | Florida, US | 1950 | II (1), III (1) | No |
| FTQY01 | Draft | II | CIP60.58 | Unknown | Unknown | Unknown | II (1), III (1) | No |
| CBYD01 | Draft | II | PW2806 | Human patient | Hong Kong | 2012 | I (1) | No |
| CBYE01 | Draft | II | PW2809 | Human patient | Hong Kong | 2012 | I (1) | No |
| ERS605480<sup>c</sup> | Draft | II | NCTC10588 | Human patient | US | 1959 | I (1), II (1) | No |
| FTQZ01 | Draft | II | CIP111067 | Unknown | Unknown | Unknown | No ICE | Yes (37) |
| CP014805.2 | Complete | III | CSID 3015183678 | Human patient | Wisconsin, US | 2016 | I (1) | No |
| ERS1197911<sup>c</sup> | Draft | III | 37-75/CIP79.29 | Human patient | St Nazaire, France | 1979 | I (1) | No |
| CP023401.1 | Complete | III | R26 | Mosquito *An.gambiae* | Stockholm, Sweden | 2005 | III (2) | No |
| CP023402.1 | Complete | III | Ag1 | Mosquito *An.gambiae* | New Mexico, US | 2012 | III (2) | No |
| CP023404.1 | Complete | III | AR4-6 | Mosquito *An.sinensis* | Sichuan, China | 2015 | III (2) | No |
| CP023403.1 | Complete | III | AR6-8 | Mosquito *An.sinensis* | Sichuan, China | 2015 | III (2) | No |
| LFKT01 | Draft | III | As1 | Mosquito *An. stephensi* | Pennsylvania, US | 2015 | III (2) | No |
| CP007547.1 | Complete | I | NUHP1 | Human patient | Singapore | 2012 | I (1), II (1), III (4) | No |
| ASYJ01 | Draft | I | NUH6 | Human patient | Singapore | 2012 | I (1), III (3) | No |
| ASYK01 | Draft | I | NUH11 | Human patient | Singapore | 2012 | I (1), II (1), III (1) | No |
| CCAC01 | Draft | I | Po0527107 | Human patient | Central African Republic | 2006 | III (1) | Yes (21) |
| CCAB01 | Draft | I | V0378064 | Human patient | Central African Republic | 2011 | III (1) | Yes (23) |
| FTPG01 | Draft | I | LDVH-AR107 | Common carp *Cyprinus carpio* | Montpellier, France | 2004 | III (1) | Yes (42) |
| CP006576.1 | Complete | I | FMS-007 | Human patient | China | 2015 | II (1) | Yes (15) |
| LWDS01 | Draft | I | EM361-97 | Human patient | Taiwan | 2000s | III (1) | No |
| AVCQ01 | Draft | I | 502 | Human patient | Birmingham, UK | 2012 | II (1) | No |
| CP016374.1 | Complete | I | F3201 | Human patient | Kuwait | 1982 | II (1) | No |
| CP016372.1 | Complete | I | JM-87 | Corn *Zea mays* | Alabama, US | 2011 | II (1), III (1) | No |
| ERS1197907<sup>c</sup> | Draft | I | 8707/CIP78.9 | Human patient | NY, US | 1962 | No ICE | Yes |
| JNCG01 | Draft | I | B2D | Human patient | Malaysia | 2013 | No ICE | No |
| FTRB01 | Draft | I | CIP111046 | Human patient | Unknown | Unknown | II (1) | Yes (6) |
| CBYF01 | Draft | I | PW2010 | Human patient | Hong Kong | 2012 | No ICE | Yes (27) |
| DRS013860 | Draft | I | GTC_10754 | Unknown | Japan | 2014 | No ICE | Yes (32) |
| LPXG01 | Draft | I | 12012-2 PRCM | Human patient | Fujian, China | 2009 | II (1), III (1) | No |

<sup>a</sup>n: number of the ICEs in the type. <sup>b</sup>n: number of the spacers in the CRISPR locus. <sup>c</sup> There are no assemblies in the NCBI database for these four strains. We assembled the genomes from Illumina reads directly.

**Figure 1.**
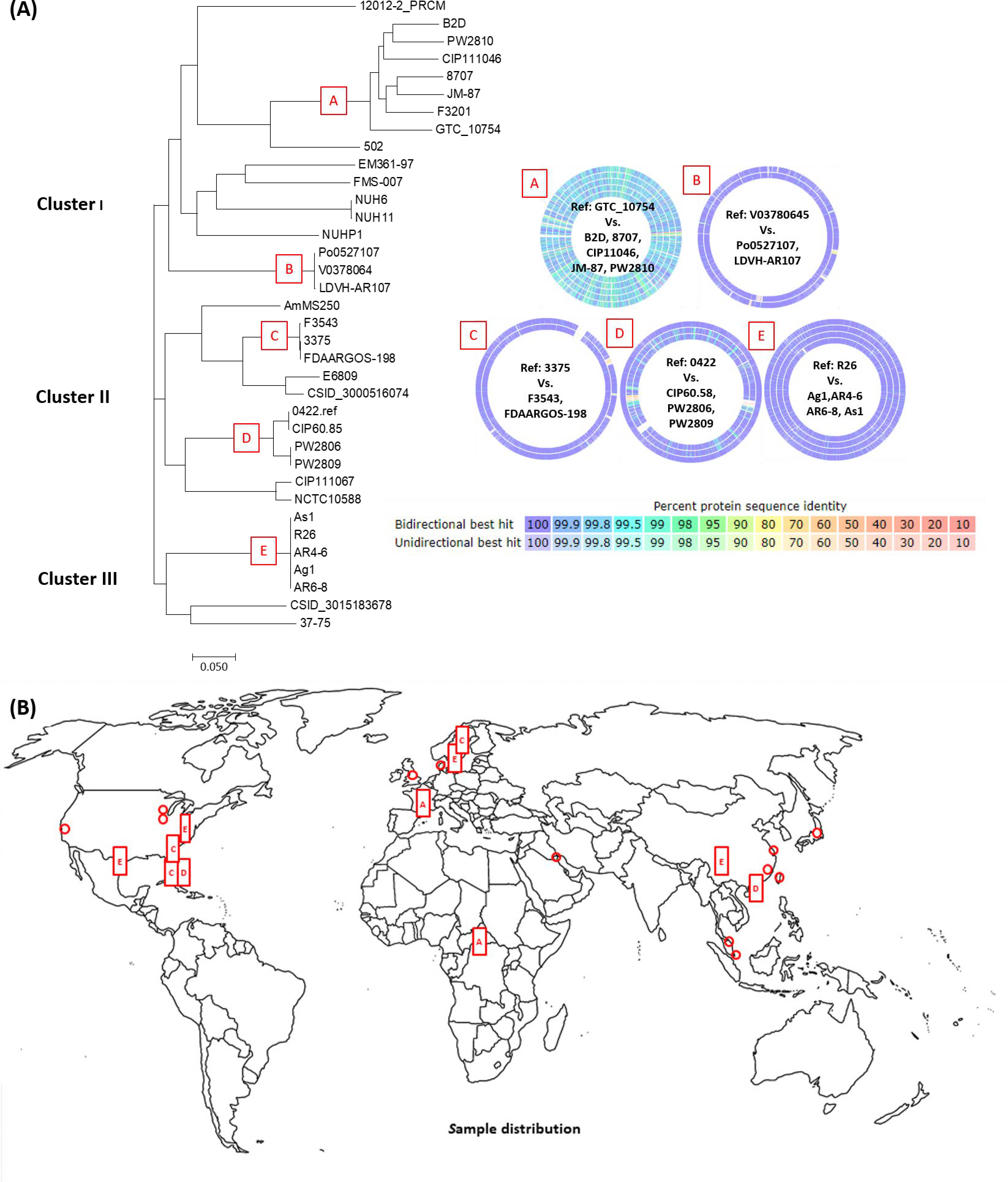
Evolutionary relationship and geographic locations of the strains. (A) The phylogenetic was tree derived from the core genome SNP comparison. Circles A-E demonstrate the protein identity between the genomes in the corresponding clades. The colour represents the percent identity when a genome was compared to the reference genome. (B) Geographic distribution of the strains. The letters correspond to the clades in part A of this figure.

### Core genome based phylogeny of the strains

The pan-genome of a bacterial species is comprised of a core and an accessory genome. The core genome is a set of genes that are conserved and shared by all strains while the accessory gene set is shared by only some strains (46, 47). Because the core genome contains essential genes and their inheritance is necessitated, they carry more reliable evolutionary information for inferring phylogeny than do accessory genes. Hence, single nucleotide polymorphism (SNP) typing based on core genome comparison has become a recognized method for accurate evolutionary reconstructions (48, 49). We used the core genome alignment to generate a SNP tree to infer phylogeny of the strains, which was implemented by Parsnp in the Harvest suit (49). Parsnp recognized 37781 anchors and 985 maximal unique matches (MUMs) in the 36 genomes, a total of 37764 anchors and MUMs were used to anchor the genome alignment and make a SNP tree (49). The tree topology appeared similar when each of the complete genomes R26^T^, 0422, CSID3015183678 and NUHP1 was used as reference for core genome alignment (data not shown). The strains were sorted into three clusters (Figure 1, Table 1). Cluster I contained 14 strains with the genome of 12012-2_PRCM at the basal position. Cluster II consisted of two clades, each included six strains. The strains in this cluster were globally distributed and collected over decades between the 1950s and the present. Cluster III had two clades as well. One was comprised of all five mosquito-derived strains, the other contained strain 37-75 obtained from a human infection in France in 1979 and strain CSID3015183678, which was a representative of the strains associated with the Wisconsin outbreak in 2016 (40). There was no indication of geography-based clustering. In each cluster, there were strains that were very closely related, which were marked as B, C, D and E in the tree (Figure 1). The clustering of these strains was corroborated by the higher percent identity in the protein sequences between the genomes. The overall identity was greater than 99.9% in the clades B, C, D and E (Figure 1). In the clade A, the strain isolated from a carp in France was closely related to the two isolates that were derived from human patients in Central Africa in 2006 and 2011, respectively (9, 39). In the clade C, 3375 was isolated from South Carolina, US, in 1957, and F3543 was isolated from Florida, US, in 1982, while FDAARGOS_198 was obtained in Sweden. In the clade D, 0422 was collected in Florida, US, in 1950, and PW2806 and PW2809 were isolated in Hong Kong in 2012. The clade E grouped the strains from three different mosquito species, *An. gambiae, An. stephensi* and *An. sinensis*, and these strains were collected in different geographic locations in separate times. The clade A presented a group of strains with certain distance, in which the identity of protein sequences was between 95-100% in a pairwise comparison. Overall, it appears that closely related strains can be widely separated both temporally and spatially.

### Identification of ICEs in the strains

Structurally, ICEs consist of the modules for genome integration and excision, conjugative transfer, and maintenance (31). We define a genome neighborhood as an ICE if it harbors genes encoding a relaxase, a coupling protein VirD4 ATPase (T4CP) and several *Tra* proteins. Among the 36 strains, 31 contained at least one ICE, and a total of 52 ICEs were detected (Table 2). According to the architecture of the modular genes, these ICEs were classified into three types (Figure 2).

**Table 2.**
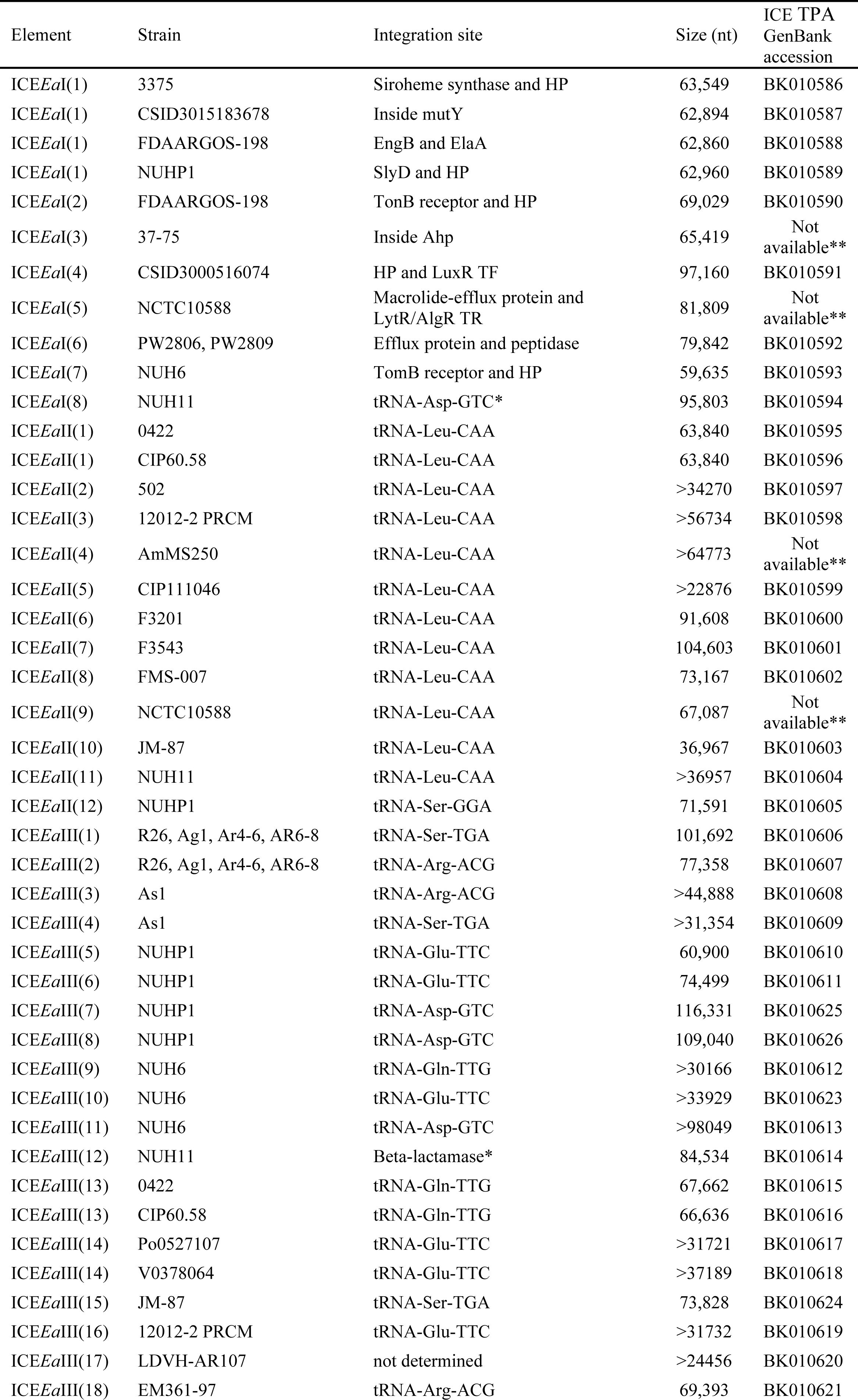
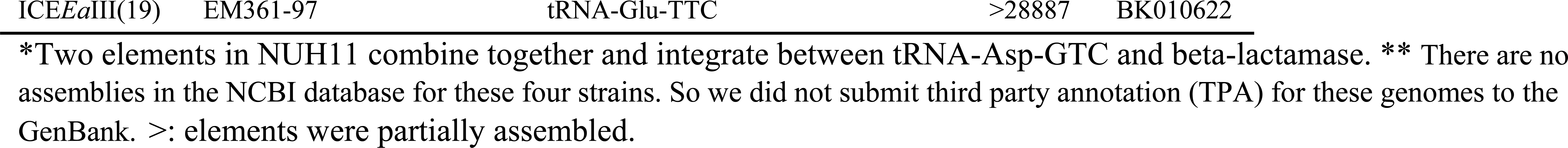
Types of ICEs identified in the strains

**Figure 2.**
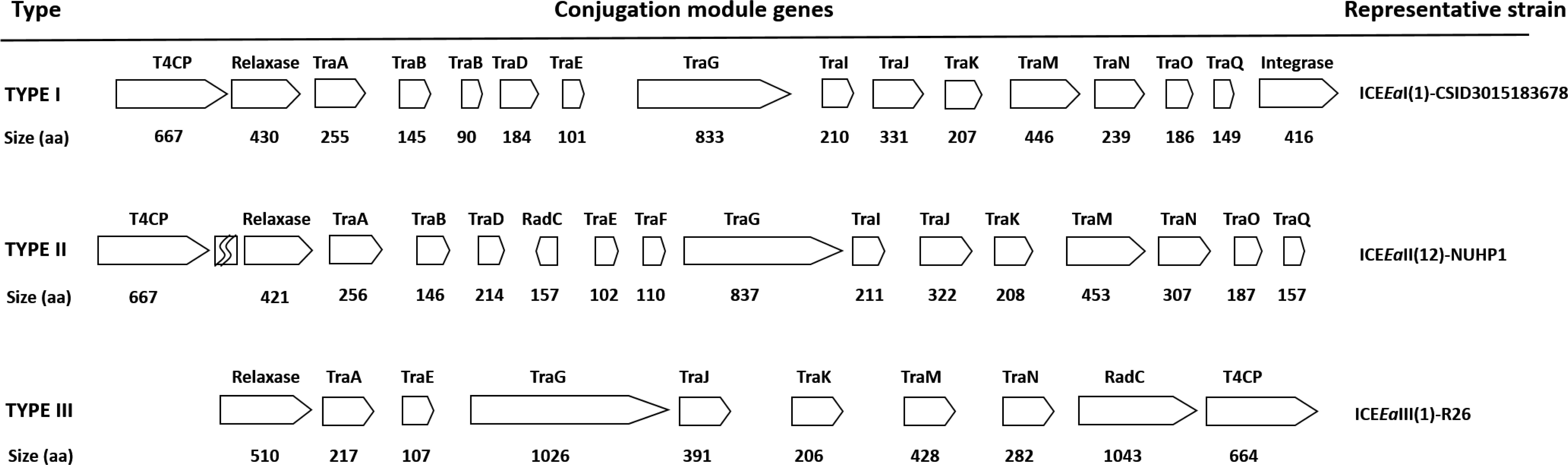
Schematic view of the architecture of conjugation modular genes in the three types of ICEs.

Type I ICEs are featured by 13 *tra* genes (*traABBDEGIJKMNOQ*) in addition to genes coding for T4CP, relaxsase and integrase. In most cases, genes *t4cp* and *relaxase* were located tandemly at upstream of the *tra* gene cassette, and *integrase* was located at downstream of the *tra* cassette (Figure 2). A total of 12 ICE*Ea*I elements were found in 11 strains, with strain FDAARGOS-198 harboring two ICE*Ea*Is (Table 2). The integration site varied. Each of the 12 elements had a distinct insertion site (Figure 3). The type I ICE in CSID3015183678 has been characterized previously, named as ICE*Ea*1 (40). This element inserted into and disrupted the gene encoding MutY, which is an adenine DNA glycosylase that is required for fixing G-A mis-pairs, making the strain more prone to mutation (40). The type I ICE in strain 37-75 inserted into the gene encoding an alkyl hydroperoxide reductase (Ahp) between codons K207 and I208. This protein is a primary scavenger of H_2_O_2_ in *E.coli* (50). Its disruption by the ICE may result in a loss of function and make the strain vulnerable to oxidative stress. The integration sites of the remaining nine type I ICEs were in the intergenic regions. The locations of these elements were listed in Table 2 and marked in Figure 3. Interestingly, The ICE*Ea*I(1) was shared by four strains, CSID3015183678 (Wisconsin, US, 2015), NUPH1 (Singapore, 2012), 3375 (South Carolina, US, 1957) and FDAARGOS-198 (Sweden, collection time unknown). The sequences of the element were nearly identical, but inwertion sites were different, indicating that the integrations occurred independently.

**Figure 3.**
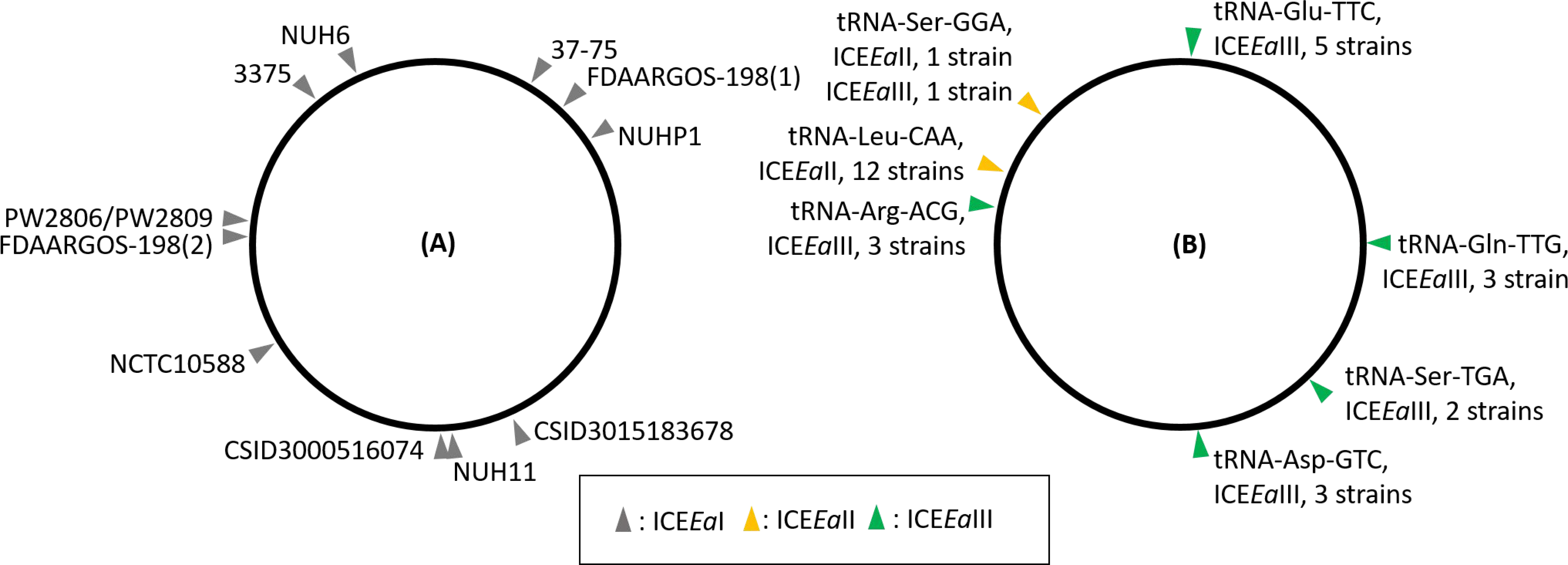
Integration sites of ICEs in different strains. (A) Location of the ICE*Ea*I integration sites. (B) Location of the tRNA genes where ICE*Ea*II and ICE*Ea*III integrated. The ICE types were color-coded. Refer to Table 2 for strain information.

Type II ICEs were identified in 13 strains. Unlike in type I, genes *t4cp* and *relaxase* were separated by one or more CDS, followed by 13 *tra* genes, *traABDEFGIJKMNOQ*. A gene is located between *traD* and *traE*, encoding a RadC-domain-containing protein (Figure 2). All 13 ICE*Ea*IIs integrated next to a tRNA gene, 12 elements reside at the 3’ end of the tRNA-Leu-CAA. Only the ICE*Ea*II in NUHP1 was inserted after the tRNA-Ser-GGA (Table 2, Figure 3). Integrases were found in 4 of the 13 elements (see below).

Type III ICEs were recognized in 16 strains. The structure was quite different from that in type I and II ICEs, with only seven *tra* genes present, *traAEGJKMN.* The *relaxase* and *t4cp* flank the *tra* genes. In 11 out of 16 elements, a gene is present after the *traN*, encoding a large protein (791-1177 aa) with a RadC domain (Figure 2). Like ICE*Ea*II, the type III ICEs after a tRNA gene. The tRNA-Arg-ACG, tRNA-Gln-TTG, tRNA-Asp-TGA, tRNA-Ser-TGA, and tRNA-Glu-TTC were targeted by ICE*Ea*IIIs (Figure 3). In NUHP1, two type III elements, ICE*Ea*III(7) and ICE*Ea*II(8), are co-located between two tRNA-Asp-GTC, and the two elements are separated by a tRNA-Asp-GTC. In NUH11, ICE*Ea*I(8) and ICE*Ea*III(12) combined as one segment, co-localized between the tRNA-Asp-GTC and the gene encoding a beta-lactamase. In most type III elements, at least one integrase gene is present.

In sum, a total of 52 ICEs were identified in 31 genomes, and 15 elements in 11 draft genomes were partially assembled. The size of ICEs is up to 116.3 kb in complete genomes. Annotated ICEs were listed in Table S1, which included predicted CDS and gene functions. Some strains had more than one ICE. No ICEs were found in five strains: PW2810, B2D, 8707, CIP11067 and GTC_10754 (Table 2).

### Phylogenetic relationship of the ICEs

To track the evolutionary history of these ICEs, the nucleotide sequences of the genes encoding relaxase, T4CP, TraG and TraJ from each ICE were compared. As shown in Figure 4, in all four gene trees, the ICEs were separated based on their types. The type I and II elements are more closely related, forming a clade distinct from the type III clade. This pattern was consistent with the type classification based on the structure of the conjugation module (Figure 2). The distribution of the ICE types did not imply geographic or temporal patterns.

**Figure 4.**
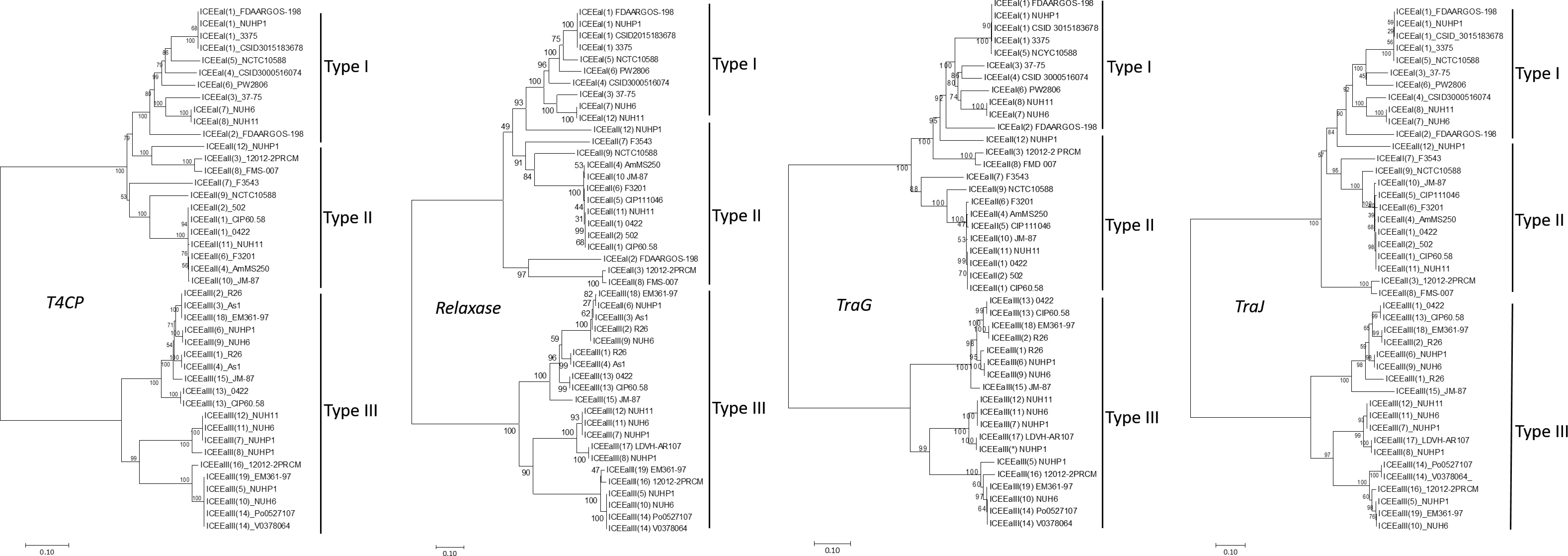
Phylogenetic relationship of the genes *T4CP, relaxase, TraG* and *TraJ*. The nucleotide sequences from different ICEs were aligned and the evolutionary history was inferred using the Neighbor-Joining method. The evolutionary distances were computed using different models to reconstruct the phylogenetic trees with the bootstrap test using 1000 replicates, which generated similar tree topology. The consensus trees generated using Kimura 2-parameter model were presented. The bootstrap values were shown on the node.

### Integrases

Integrases, required for ICE integration and excision, could be tyrosine or serine recombinases, or DDE transposases containing an acidic amino acid triad (DDE)(32). Tyrosine recombinases were found in all type I and III ICEs, and DDE transposases were identified in four type III ICEs. In the 12 type II ICEs, only four elements carried integrases. Two serine recombinases were detected in NCTC10588 and one was identified in JM-87. A DDE transposase was found in FMS-007, and a tyrosine recombinase was detected in NUHP1. The remaining 9 type II ICEs lack co-localized integrases (Table S1).

### Cargo genes

Restriction and modification (R-M) systems provide an innate defense against invading DNA to protect the genome stability. Mobile genetic elements (including ICEs) usually carry an R-M system, which enables an evolutionary interplay between the mobile genetic elements (MGEs) and their hosts (51). The DNA modification methylase, type I R-M system, and type II restriction enzymes were prevalent in the ICEs of all three types. In addition, the anti-restriction protein ArdA was detected in three type II ICEs and one type III ICE. Some ICEs also carried DNA topoisomerase, DNA helicase, DNA primase, DNA polymerase. These enzymes may be involved in the integration process, and perhaps in DNA replication while the ICE is in its plasmid state. Genes encoding proteins in the resistance-nodulation-division (RND) family (52), a multidrug efflux pump system found in Gram-negative bacteria, are prevalent in the ICEs. The genes coding for three key component proteins of the tripartite RND complex (the outer membrane protein, membrane fusion protein and inner membrane protein), are carried in the type I ICEs in six strains, one of the three type III ICEs in NUHP1 and the type II ICE in F3543 (Table S1). ABC transporters for various substrates, such as manganese, potassium and oligopeptides were found in all ICEs. TonB-dependent receptors for siderophore import or carbohydrates uptake (SusC) are present in some ICEs. There are various transcriptional regulators in the ICEs, such as AraC family, ArsR family, MarR family, HxlR family, and TetR family. The AraC family transcriptional regulators (AFTRs) are most prevalent, a total of 36 these were found in 19 strains. In addition, two-component regulatory system are present in the type I ICEs in three strains.

### CRISPR-Cas loci

The CRISPR-Cas system is a prokaryotic adaptive defense machinery against invading nucleic acids (53, 54). The type II-C CRISPR-Cas system was detected in nine strains (Table 1), which is featured by a CRISPR array and the genes encoding Cas9, Cas1 and Cas2 proteins. In these nine strains, the CRISPR-Cas system is located downstream of the gene encoding a cobalt-zinc-cadmium resistance protein CzcD. In a CRISPR array, the direct repeat is 47 nt, and the spacer is 30 nt in length. The number of spacers varies between 6 in strain CIP111046 to 47 in strain CIP111067. No CRISPR locus was detected in the draft assembly of strain 8707, but one complete direct repeat was identified in the assembly. Likely the CRISPR locus was not assembled well in the draft genome. The CRISPR loci of strains Po0527107 and V0378064 have been described previously (39). Po0527107 has 21 spacers and V0378064 has the same 21 plus two more spacers. Both strains were isolated in Bangui, the Central Africa Republic, but five years apart (9, 55). Strain LDVH-AR107 has two CRISPR loci, each with 21 spacers. LDVH-AR107 shares 12 spacers with both Po0527107 and V0378064 despite being isolated on a different continent, from the internal organ of a common carp *Cyprinus carpio* collected in 2004 in Montpellier, France (Table 1).

### Discussion

Many genomes have been sequenced for *E. anophelis* strains derived from different sources including mosquitoes, human infections, hospital environments, fish and corn stems. This genome availability enabled a comparative genomics approach to investigate the genetic architecture and repertoire of the *E. anophelis* population.

### Core genome based phylogeny

By definition, the core genome is shared by all strains and consists of essential genes that are vertically transmitted while genes in the accessory genome are present only in a subset of strains. The sum of the core and accessory genomes in all strains constitutes a pan-genome for the species (47, 56, 57). Due to its inheritability, the core genome is intrinsically suitable for inferring phylogeny (49). The core genome phylogenetic analysis separated the strains into three clusters, however, genetically similar strains do not show a spatial or temporal aggregation pattern. As an example, clade C (Figure 1) contains closely related strains F3543, 3375 and FDAARGOS-198, the locations where they were isolated were Florida in 1982, South Carolina in 1957 and Sweden (collection time unknown), respectively (Table1, Figure 1). On the other hand, the mosquito-associated strains were very closely related, and distinct from the strains derived from human infections (Figure 1), despite having been isolated from three different mosquito species, and from both lab reared (19, 21) and wild caught mosquitoes (20). The possibility that they might be lab contaminants was carefully evaluated, and rejected based on the timeline of sample collection, processing and sequencing. Furthermore, the presence of *E. anophelis* in the gut microbiota of wild populations of *An. gambiae* in Kenya and *An. sinensis* mosquitos in Shandong, China, has been confirmed from the shotgun metagenomic sequencing data (Xu, unpublished data). Association of *E. anophelis* has been documented in several different mosquito species, including *An. gambiae* (58), *Culex quinquefasicatus* (59) and *Aedes aegypti* (60), and a role for the bacteria in larval development has been hypothesized. Taken together, these results indicate that there is selection in the mosquito gut environment for these specific strains.

### ICEs in the strains

Genome diversification promotes bacterial adaptation and evolution. ICE, as a type of mobile genetic elements, contributes significantly to the pan-genome reservoir with potentially adaptive genes (28, 31). Several ICEs have been found in the taxa of *Bacteroides* (29, 61-64), which belong to the family Bacteroidaceae in phylum Bacteroidetes. Of these elements, CTnDOT in *B. thetaiotaomicron* has been studied extensively (65-68). The ICEs identified in *E. anophelis* are structurally distinct from CTnDOT.

The conjugation machinery is a necessary component of ICEs. Relaxase, coupling protein and *tra* proteins are required for element excision from the genome and transfer to a recipient (69). Conserved features indicative of conjugative elements have been used for identifying ICEs in a wide variety of genomes (29). Using this approach, we categorized the ICEs in *E. anophelis* strains into three types based on the architecture of the genes in the conjugation module and the phylogeny of sequences from four genes in the module (Figure 2, 4). Type II and III ICEs tended to integrate adjacent to a tRNA gene, while the type I ICEs used a non-tRNA gene region for integration. In some strains, the type I ICE insertion into a gene resulted in the loss of function. For example, in CSID3015183678 the integration of ICE disrupted the gene *mutY*, which is required for excision of G-A mismatch (70, 71). The disruption of *mutY* by ICE*Ea*I resulted in a higher rate of nonsynonymous to synonymous substitution in the strain and subsequent sublineages, which may fuel the adaptive capacity of an outbreak strain (40). In strain 37-75, the insertion of a type I ICE inactivates the gene encoding alkyl hydroperoxide reductase (Ahp). As the Ahp protein is a scavenger of hydrogen peroxide in *E. coli* (50), the disruption of the *ahp* gene by ICE may affect host defense against elevated oxidative stress. Twelve type II ICEs used tRNA-Leu-CAA as the integration locus, however, these elements lacked an integrase. This warrants further investigation to elucidate whether or not the type II ICEs are mobile without an integrase in the element. The presence of diverse integration loci, including tRNA gene and non-tRNA loci (Figure 3), suggests a high degree of genome plasticity of the *E. anophelis* in favor of a dynamic genetic repertoire of the pan-genome accommodating mobile elements like ICEs (28, 63).

Among the cargo genes carried by the ICEs for which a function could be identified, most are involved with host defense, nutrient acquisition, or transcription regulation. Innate immune mechanisms like R-M systems provide host defense by protecting against invading DNA (51, 72), and the RND type multidrug efflux pumps enable the chemical defense by recognizing and expelling various structurally diverse compounds and toxins (73), including antibiotics, as shown previously in R26^T^ (37), and the strains associated with the outbreak in Singapore (35, 36). Regarding nutrient acquisition, a variety of ABC transporters and TonB dependent receptors are present in the ICEs, which greatly enhances the host capability to satisfy the nutrient needs in the environment they thrive. The ICEs also equip the host with a variety of transcriptional regulators, with those in the AraC family being particularly prevalent (Table S1). These transcription regulators can sense various chemical signals involved in carbon metabolism, quorum-sensing signaling, virulence and stress response, which leads to the expression of a global gene repertoire (74-77). Such genetic capability brought in by ICEs enables the host strains a broader adaptive flexibility to eco-niche changes, which may contribute to establishment of an infection in humans as well.

### CRISPR loci

CRISPR-Cas system is an adaptive immune mechanism against invading nucleic acids. Spacer sequences are acquired and integrated into the CRISPR locus. Upon encountering the same non-self invaders, the spacers will be transcribed into guide RNAs to direct Cas nuclease to cleave specific target sequences (53, 78). In this study, the class II CRISPR-Cas system was identified in nine strains of *E. anophelis.* A total of 209 spacers were recognized in eight strains. These spacers reflected previous events of the horizontal transfer of mobile elements or phage infections these strains have experienced in the past. Indeed, it has been shown by Breurec *et al.* (39) that strain V0378064 (isolate E18064 in (39)) obtained in 2011 contained two newly acquired spacers, S22 and S23, which are a perfect match to two protospacers from the strain Po0527107 (isolate E27107) isolated in 2006. In fact, the two protospacers reside in a phage-derived mobile genetic element integrated at the 3’ end of tRNA-Arg-ACG in Po0527107. The phage element is absent from V0378064, likely due to the action of the CRISPR-Cas mediated defense.

Of nine strains that carry a CRISPR-Cas system, five have ICEs as well. The co-existence of mobile elements and CRISPR defense system suggests an evolutionary balance between the genome stability and plasticity. In the family *Flavobacteriaceae*, the CRISPR-Cas systems in *Flavobacterium columnare* were shown to play a role in host-phage interactions, which drives long-term genome coevolution through arms-race-like mechanisms between the host and phages (79), serving as an example of genome stability. On the other hand, the CRISPR-Cas machinery appeared not to play a role as a resistance mechanism against phages in fish pathogenic *Flavobacterium psychrophilum* (80). Availability of the *E. anophelis* strains that possess CRISPR-Cas with and without ICEs enables further studies on co-evolution of mobile elements and host defense systems in the bacteria.

### Conclusion

In this study, we identified ICEs in the genomes of 36 *E. anophelis* strains isolated from a wide array of hosts, collected at diverse geographic locations over several decades. The ICEs can be categorized into three types based on the distinctive architecture of conjugation module and integration sites. The identification of the ICEs enables further studies on the genetic diversity of the pan-genome and its impact on the virulence of this global opportunistic pathogen.

## Materials and Methods

### Bacterial strains

In this study, complete genomes of 13 strains and draft genomes of 23 strains were analyzed (Table 1). Mosquito-derived strains were isolated from *An. gambiae* (R26^T^, Ag1), *An. stephensi* (As1) and *An. sinensis* (AR4-6, AR6-8). Strain CSID3015183678 was one of the strains associated with the Wisconsin outbreak 2015-2016, which has been characterized in (40), and strain NUHP1 was a representative of the strains isolated in the Singapore outbreak, the genomes of these strains have been described (35). Strain LDVH-AR107 was derived from a common carp *Cyprinus carpio*. Strain JM-87 was isolated from maize *Zea mays,* which was originally described as *E. endophytica*. The genome comparison put it as a synonym of *E. anophelis* (7). The other strains were all derived from human patients or hospital environments, collected from 1950 to 2016.

The draft genomes of strains LDVH-AR107, 8707, NCTC10588, AmMS250 and 37-75 were reassembled using Illumina reads downloaded from SRA database. The reads were *de novo* assembled by CLC genomics workbench v 10.1.1. The genomes were annotated using the SEED and Rapid Annotations using Subsystems Technology (RAST) at the RAST server (81).

### Phylogenetic relationship of the strains based on core genome comparison

The phylogenomic relationships of the strains were estimated by the Harvest suite (49). The core genome that is shared by all strains was identified by multiple genome alignment, and SNPs in the core genome were typed to infer phylogenies between the strains implemented by Parsnp module in the Harvest suite (49). Four complete genomes R26^T^, 0422, CSID3015183678 and NUHP1, each was used iteratively as the reference for tree construction, and tree topology of all four core genome trees was unaffected by choice of reference genome. The tree with 0422 as the reference was depicted in Figure 1.

### Identification of ICEs

Pairwise genome comparison based on protein identity was used to identify variable regions. To identify an ICE, the genome was searched for a cluster of genes coding for a relaxase, a coupling ATPase (T4CP) and transfer (Tra) proteins including a VirB4 ATPase (TraG) in the conjugation module. These proteins are the key components of an ICE (28, 69). The boundary of an element was delimited as between the two open reading frames (ORFs) that flank the ICE. The genome of the type strain R26^T^ was used as a reference to mark the integration sites. Each ICE was categorized into one of the three Types based on its gene structure (Figure 2). The ICEs were named based mainly on the nomenclature proposed by Burrus *et al.* (82): the acronym ICE was followed by the initials of the name of the bacterium (*Ea*), a Roman numeral as type, a strain name and a ordinal number in brackets to identify the same sequence if it is encountered in a different strain. For example, the type I ICE found in both NUHP1 and CSID3015183678 would be named ICE*Ea*I(1)_CSID3015183678 and ICE*Ea*I(1)_NUHP1, respectively. Each type has its set of numbers; for example, the four type III ICEs in NUHP1 are designated as ICE*Ea*III(1)_NUHP1 thru ICE*Ea*III(4)_NUHP1.

## GenBank accession numbers

Accession numbers of the genomes used in this study were listed in Table 1. GenBank accession numbers for ICE sequences were listed in Table 2.

## Supplemental Material

Table S1. Annotated ICEs in the strains. (excel file)

## Author contributions

JX conceived and designed study. JX, DP, YL,QX collected genome data and performed data analysis. JX and AN wrote the manuscript.

## Acknowledgements

This work was supported by the National Institutes of Health [SC1AI112786 to J.X.] and the National Science Foundation [No. 1633330 to J.X.] and CDC program funds designated for the study of emerging infectious agents. The content is solely the responsibility of the authors and does not necessarily represent the official views of the National Institutes of Health and the National Science Foundation, and the Centers for Disease Control and Prevention. Mention of company names or products does not constitute endorsement. Funding for open access charge: National Institutes of Health [SC1AI112786].

